# EasyCAPS: A web tool for restriction-based genotyping and rational CRISPR-Cas9 donor design

**DOI:** 10.64898/2026.04.17.719238

**Authors:** Lucas Souza de Bem, Jeferson Gross, Ana Paula Jacobus

## Abstract

Tracking Single Nucleotide Polymorphisms (SNPs) following CRISPR-Cas9 genome editing is a critical yet often labor-intensive step in modern genetic research. Although Sanger sequencing is the conventional method for definitive confirmation, it typically requires substantial time to generate results. In contrast, PCR-based restriction methods like CAPS (Cleaved Amplified Polymorphic Sequence) and dCAPS (derived CAPS) offer rapid and cost-effective alternatives. However, existing dCAPS primer design tools suffer from significant limitations and were largely developed for tracking polymorphisms in plant genomes. Concurrently, CRISPR-Cas9 gene editing requires strategies to prevent the re-cleavage of the edited allele, typically involving the modification of the Protospacer Adjacent Motif (PAM). To address these challenges, we developed EasyCAPS, a web-based tool that integrates dCAPS primer design with advanced functionalities for CRISPR experiments. EasyCAPS overcomes the shortcomings of previous software by enabling restriction enzyme pre-selection and optimizing designs for complex DNA sequences. Its key innovation is the “Hiding PAM” feature, which designs synonymous mutations to mask the Cas9 recognition site while accounting for codon usage bias, thereby facilitating one-step allelic exchange. The utility of the tool was demonstrated through practical applications targeting the *HTA1*, *PHO84*, and *CAT5* genes, significantly accelerating both genotyping and gene editing processes. We conclude that EasyCAPS is an accessible solution that effectively streamlines molecular biology workflows.

## Introduction

The detection of genetic variants, such as single-nucleotide polymorphisms (SNPs) and small insertions or deletions (indels), is a fundamental component of genetics studies. Although larger indels can be easily discriminated by conventional electrophoresis of PCR products, smaller variants, especially SNPs, remain indistinguishable using methods based solely on fragment size. In such cases, Sanger sequencing remains the reference method. However, its high cost and the time required for sample preparation and access to specialized equipment limit its routine application for screening large numbers of samples in many laboratories.

As a cost-effective and accessible alternative, the analysis of PCR products digested with restriction enzymes has been established as a robust strategy for genotyping. The Cleaved Amplified Polymorphic Sequence (CAPS) method (Fig. 1A) allows for allelic differentiation based on the presence or absence of natural restriction sites in the amplicons [1]. However, the reliance on the fortuitous occurrence of such sites limits the applicability of the method. To overcome this barrier, the dCAPS (*derived CAPS*) technique (Fig. 1B) utilizes primers with strategically designed mismatches to generate an artificial restriction site adjacent to the mutation, enabling genotyping in the absence of natural sites [2]. Despite their theoretical efficacy, the manual design of dCAPS primers is laborious and error-prone. Moreover, currently available tools [3,4] exhibit critical limitations, including restrictions on sequence length, low flexibility in enzyme selection, inability to integrate the simultaneous search for natural CAPS and dCAPS, and algorithms that are often ineffective in detecting all potential cleavage possibilities.

**Fig. 1.**
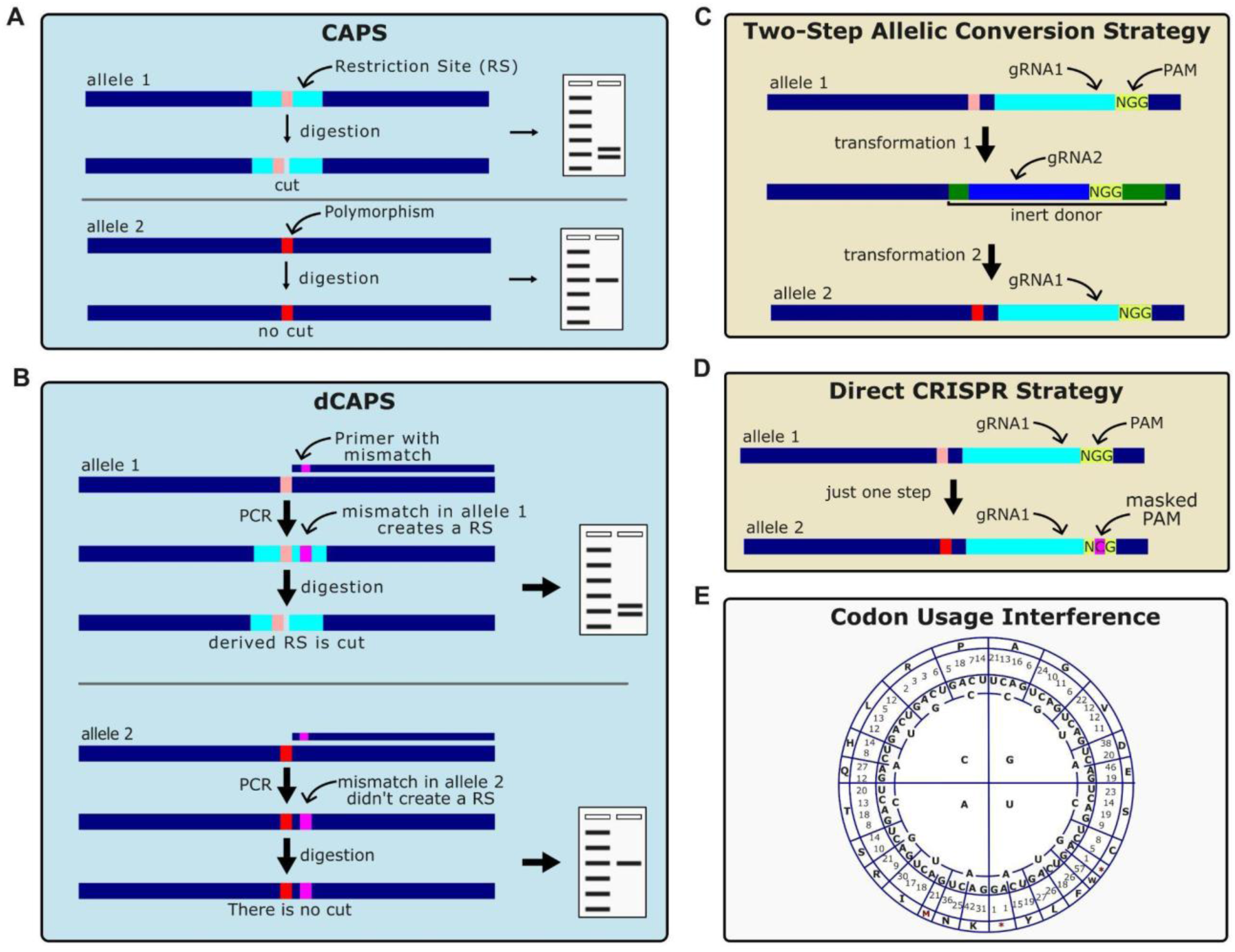
Overview of strategies for polymorphism analysis and CRISPR-mediated allelic conversion. **(A)** The CAPS method exploits a natural SNP that creates or destroys a restriction enzyme recognition site, allowing allele differentiation through digestion. **(B)** In the dCAPS strategy, a mismatch is intentionally introduced via the PCR primer to artificially generate a restriction site in the PCR product derived from only one of the alleles, enabling differentiation when no natural restriction site is present. **(C)** The conversion of alleles is performed in two sequential transformations to prevent re-cleavage of the edited allele. First, gRNA1 directs the insertion of an inert donor cassette that replaces the original target site with a new sequence that is recognized by gRNA2. In the second step, gRNA2 is used to introduce the desired variant, generating Allele 2. **(D)** In this strategy, allelic replacement is achieved in a single transformation step. The repair template carries a silent mutation within the PAM sequence. This masks the Cas9 recognition site in the newly edited allele, preventing recurrent cleavage and ensuring editing efficiency without altering the protein sequence. **(E)** The chart illustrates the redundancy of the genetic code. Although synonymous mutations (used in D) do not alter the protein sequence, the preferential use of certain codons (codon usage bias) can impact gene expression, translation speed, and protein folding.

In parallel, the widespread adoption of the CRISPR-Cas9 system for precision editing has imposed new challenges in experimental design [5]. This complexity is particularly evident when manipulating essential genes, where the conventional two-step strategy (Fig. 1C), involving gene disruption or deletion followed by correction, is infeasible because the transient inactivation of the locus would result in cell lethality. In such scenarios, direct one-step editing is mandatory. However, in direct allelic replacement strategies via homologous recombination, preserving the Protospacer Adjacent Motif (PAM) sequence in the donor DNA can induce re-cleavage of the newly edited allele, significantly reducing editing efficiency [6,7]. To facilitate direct editing and mitigate this risk, an efficient approach involves introducing silent mutations into the PAM or protospacer sequence (Fig. 1D), thereby masking the nuclease recognition site without altering the primary sequence of the target protein [8].

Nevertheless, the introduction of these “silent” mutations requires caution. Although the amino acid sequence is preserved, the choice of synonymous codons is not biologically neutral. Codon usage bias (Fig. 1E) directly influences mRNA stability, translation speed, and accuracy, as well as proper protein folding. The inadvertent substitution of a frequent codon with a rare one can create translation bottlenecks, potentially reducing gene expression or altering protein function [9]. Therefore, the design of PAM mutations must balance the abolition of the Cas9 recognition with the maintenance of the translational efficiency of the organism.

In this context, we present EasyCAPS, an interactive web tool developed to integrate genotyping and gene-editing planning. The software facilitates the automatic identification of natural and derived restriction sites (CAPS and dCAPS) and assists in the rational design of donor sequences, suggesting silent mutations in the PAM, alongside codon usage analysis. Thus, EasyCAPS provides a unified solution to enhance the precision and efficiency of both mutant screening and gene editing via CRISPR-Cas9 technology.

## Materials and methods

### Implementation and architecture

The EasyCAPS web application was developed using Python 3.14.0 as the core programming language. The backend infrastructure was built upon the Flask 3.1.0 microframework, utilizing Jinja2 3.1.6 for server-side template rendering and Bleach 6.2.0 for input sanitization and security. The web interface was designed using standard HTML5, CSS3, and JavaScript to ensure responsiveness and cross-browser compatibility without relying on heavy frontend frameworks. The development process involved the use of Large Language Models (LLMs), including ChatGPT, Gemini, and Copilot, which assisted in code syntax generation, debugging, and optimization of logic routines. The source code is maintained on GitHub (https://github.com/olucasdebem/EasyCAPS_git) and is deployed directly to the Render cloud platform, ensuring that the live web application operates as a synchronized mirror of the repository. The website is available at https://easycaps-app.onrender.com/.

### Algorithm design and logic

The core algorithms for restriction site identification and mismatch generation were designed based on the fundamental principles established by previous tools such as dCAPS Finder 2.0 [3] and indCAPS [4]. However, the logic was completely rewritten to improve flexibility. The computational pipeline of EasyCAPS was designed to process inputs sequentially through four distinct modules: Data Preprocessing and Validation, Core CAPS/dCAPS Identification, Silent Mutation Engineering (Donor/PAM), and Output Rendering.

Upon submission, input sequences (Allele 1 and Allele 2) and guide RNAs (gRNAs) undergo strict sanitization using the Bleach library to prevent injection attacks. The algorithm validates the input against IUPAC nucleotide codes and enforces a maximum length limit of 200 bp to ensure efficient processing. The sequences are standardized to uppercase, and all whitespace is removed. Simultaneously, the system generates the reverse complement (RC) sequences and performs *in silico* translation of the forward strands to establish the reading frame. A differential alignment algorithm identifies the position of SNPs or small indels, highlighting the variable regions between the two alleles.

The core analysis is performed based on a user-defined restriction enzyme library. The algorithm isolates a window of approximately 60 nucleotides (9 codons upstream and downstream) centered on the target SNP, or the most upstream SNP if multiple variants exist. The tool scans both alleles (Forward and RC) for potential restriction sites, allowing a user-defined number of mismatches from 0 to 3. To ensure assay specificity, the identified sites are filtered based on strict criteria: the SNP must be located within the enzyme’s recognition site, at the specific cut site, or between the mismatch position and the cut site. Sites that do not meet these criteria are discarded. Subsequently, the algorithm compares the restriction profiles of Allele 1 versus Allele 2, retaining only enzymes that cleave one allele exclusively. Unique sites with zero mismatches are classified as Natural CAPS. If no natural sites are found, the algorithm proceeds to dCAPS design, where mismatches are iteratively introduced into the primer sequence adjacent to the 3’ end to artificially create a restriction site that depends on the SNP. These putative dCAPS primers then undergo an *in silico* validation step to confirm that the introduced mismatch, combined with the template SNP, generates a differential restriction pattern for the target gene.

For CRISPR-Cas9 editing strategies, the tool includes a module for designing repair templates (donors) and masking the PAM sites. The algorithm first retrieves a codon usage table for the selected target organism. It then scans the donor sequence for positions where single-nucleotide substitutions can create new restriction sites (Silent CAPS) with a maximum of one mismatch, without altering the amino acid sequence. Simultaneously, for the “Hiding PAM” analysis, the tool identifies PAM sequences associated with the input gRNAs and generates variants that abolish Cas9 recognition while preserving the protein sequence. All generated synonymous mutations are evaluated against the organism’s codon usage table. The tool calculates and displays the ratio between the frequency of the original codon and the new codon, flagging rare codons to prevent negative impacts on translation efficiency.

Finally, the results are aggregated into structured objects containing natural sites, validated dCAPS primers, and engineered donor sequences. The frontend dynamically renders these results, grouping analyses and providing a visual representation of the variations, restriction enzymes, and codon usage information.

### Application of EasyCAPS for genome editing and genotyping strategies

To validate the utility of the EasyCAPS tool in practical scenarios, we performed allelic replacements in *Saccharomyces cerevisiae,* targeting the *PHO84*, *HTA1*, and *CAT5* genes [10]. Strategies for donor design, PAM masking, and genotypic validation were established using the specific software modules.

In this study, the *S. cerevisiae* strain S288C was used as the biological model. Cultures were propagated in YPS-rich medium composed of 10 g/L yeast extract, 20 g/L peptone, and 20 g/L sucrose, with the addition of 15 g/L agar for solid media. For the selection of transformants, YPS plates were amended with specific antibiotics as necessary: nourseothricin (100 µg/mL), geneticin (200 µg/mL), zeocin (250 µg/mL), or hygromycin B (300 µg/mL) [11].

Allelic replacements were generated using the EasyGuide CRISPR-Cas9 system [12]. Briefly, specific gRNAs targeting loci of interest (*PHO84*, *HTA1*, and *CAT5*) were cloned into the pEasyG3 vector. Donor DNA fragments, which serve as repair templates for homologous recombination, were generated by PCR with oligonucleotides designed via the EasyCAPS tool. These repair templates contained the desired point mutations and synonymous mutations for PAM masking or restriction site creation (CAPS/dCAPS) [13,10]. Yeast transformations were performed using the standard lithium acetate/polyethylene glycol method [14]. Successful editing events were screened by colony PCR and restriction digestion analysis (CAPS/dCAPS) and were subsequently confirmed by Sanger sequencing.

## Results

### Interface and operation of EasyCAPS

The EasyCAPS interface was designed to offer an intuitive and centralized workflow, eliminating the need to navigate multiple pages or tabs. The main dashboard is divided into three logical modules that guide the user from sequence input to the configuration of the gene editing parameters (Fig. 2). In the upper module (Fig. 2A), the researcher inputs the nucleotide sequences of the two alleles to be compared (e.g., wild-type and mutant). Crucially, for analyses involving coding regions, such as donor design or protein translation verification, the input sequences must be provided in the correct reading frame (frame 0) to ensure accurate downstream codon usage calculations. The algorithm accepts raw or formatted sequences and automatically removes spaces and non-nucleotide characters. In this panel, the user also defines the mismatch threshold allowed for dCAPS primer design, typically set to 1 to ensure PCR specificity, allowing the tool to automatically switch between searching for natural sites (CAPS) and engineering artificial sites (dCAPS) as needed.

**Fig. 2.**
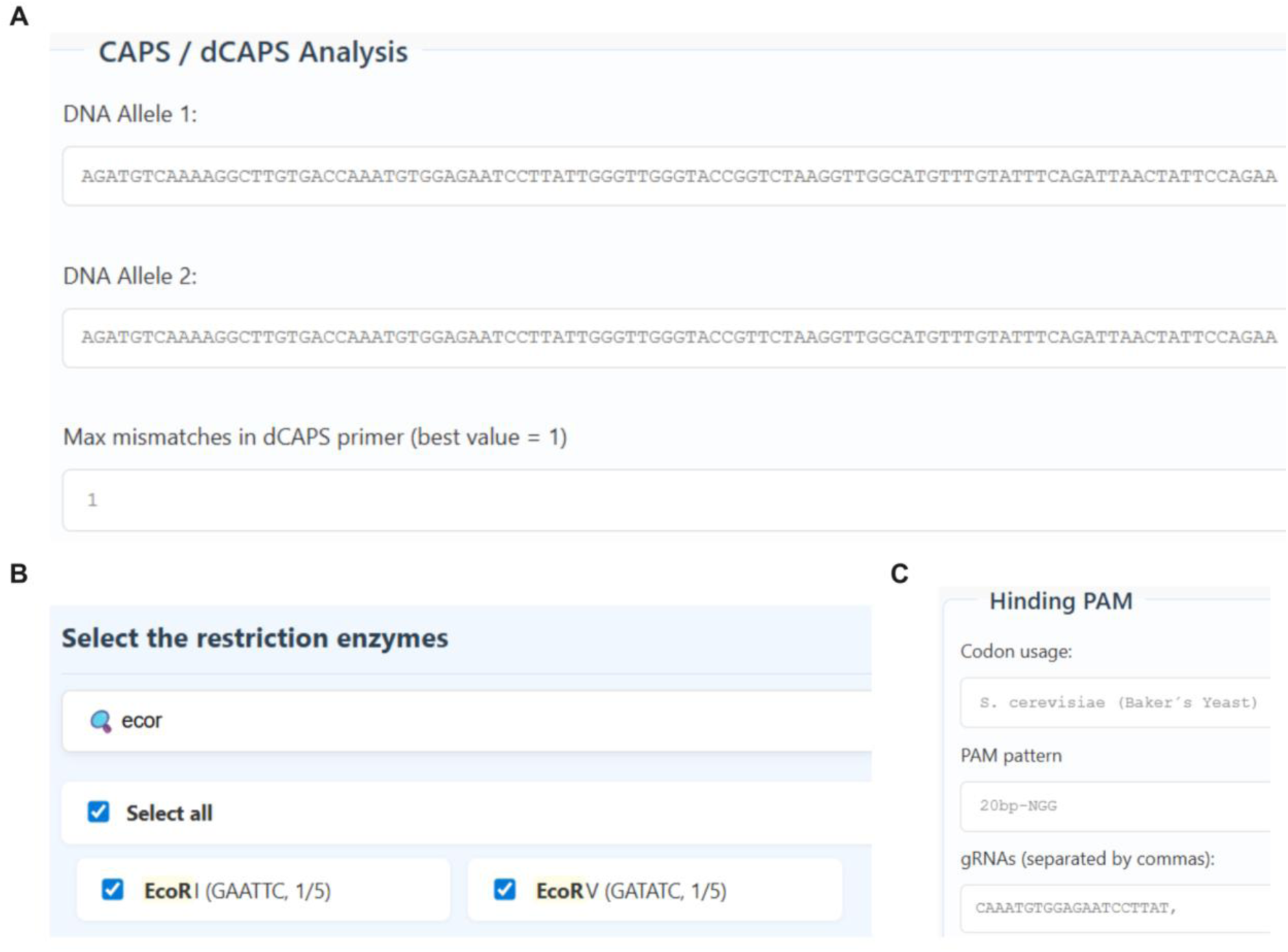
EasyCAPS web interface and input modules. **(A)** The dashboard allows users to input the nucleotide sequences for both alleles (e.g., Wild-Type and Mutant) and define the threshold for maximum mismatches allowed in dCAPS primer design (typically set to 1). **(B)** This module provides a flexible filtering system for the restriction enzyme library. Users can search for enzymes by name or recognition sequence and choose to analyze a specific subset or the entire database using the “Select all” option. **(C)** Integration of gene editing parameters for silent mutation strategies. Users select the target model organism to load the appropriate codon usage table, define the required PAM motif (e.g., NGG), and input the gRNA sequences to enable PAM masking and synonymous mutation analysis.

The second module (Fig. 2B) offers full control over the restriction enzyme library. Unlike previous tools that limit the user to pre-defined lists [3,4], EasyCAPS allows for real-time searching by enzyme name or recognition sequence. The user can choose to analyze all available enzymes (“Select all”) for an exploratory scan or select a specific subset of enzymes available in their laboratory, optimizing experimental cost and logistics.

Finally, the lower module (Fig. 2C) is optional and integrates unique gene editing functionalities. For strategies requiring silent mutations, such as Hiding PAM or creating Silent CAPS in donors, the user selects the target organism from a dropdown menu (e.g., *S. cerevisiae*, *E. coli*, *H. sapiens*), which automatically loads the appropriate codon usage table. The user then inputs the PAM pattern (default NGG for Cas9) and the list of up to 10 gRNA sequences, typically obtained from specialized design tools like CRISPOR [15]. This integration enables the algorithm to evaluate not only the creation of restriction sites but also the biological impact of suggested mutations on translational efficiency.

### Output visualization and CRISPR mutagenesis strategies

Upon execution, the EasyCAPS results interface presents a comprehensive alignment of the input sequences (Fig. 3A), where the target polymorphism is visually highlighted in yellow. For genotyping, the tool outputs distinct panels for each allele. It prioritizes the identification of natural restriction sites (CAPS), displaying the enzyme name, recognition pattern, and specific cut site (Fig. 3B and 3D). Beyond the natural sites, the software generates dCAPS strategies, providing sequences for forward or reverse primers containing the necessary mismatches (highlighted in red) to create artificial restriction sites, as demonstrated by the *Acc*I and *Acl*I examples (Fig. 3C and 3E).

**Fig. 3.**
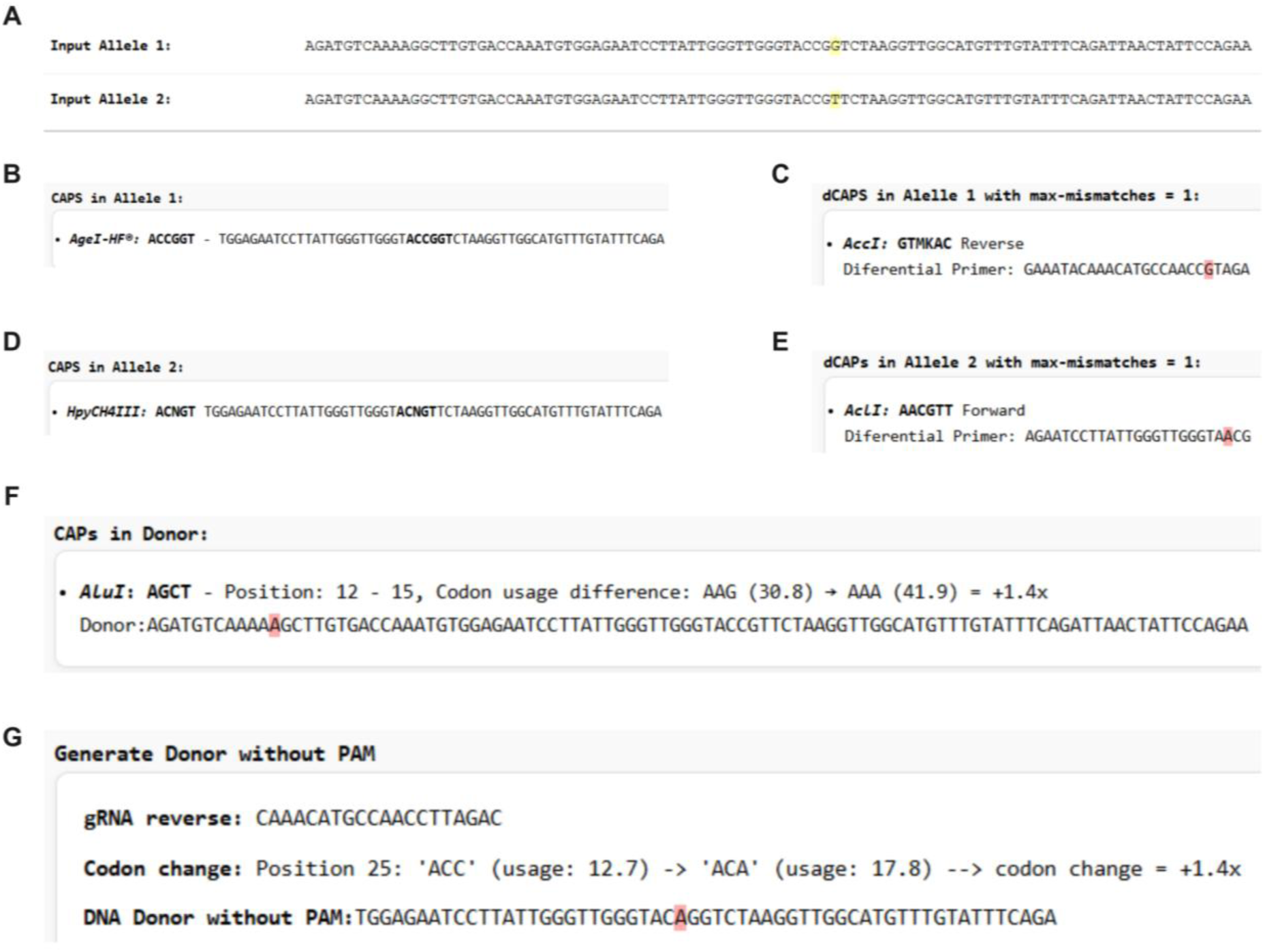
EasyCAPS Output and Analysis Modules. **(A)** Sequence alignment panel highlighting the target SNP (yellow) or variable region. **(B, D)** Identification of natural CAPS markers (bold) for Allele 1 and Allele 2, respectively. **(C, E)** Design of dCAPS primers with mismatched bases (red) to create artificial restriction sites. **(F)** Engineering of silent restriction sites within the donor sequence, displaying the created restriction site (*Alu*I) and the codon usage fold-change (+1.4x). **(G)** PAM masking module (“Hiding PAM”), showing the introduction of synonymous mutations to abolish Cas9 recognition while maintaining favorable codon usage statistics.

Beyond standard genotyping, EasyCAPS includes specialized functionalities for rational donor design in mutagenesis and CRISPR-Cas9 experiments. The “CAPS in Donor” module (Fig. 3F) automates the engineering of silent sites by scanning the donor sequence to identify positions where single-nucleotide substitutions can introduce a novel restriction site to serve as a genetic marker. These mutations are designed to be synonymous, preserving the amino acid sequence. To ensure biological viability, the tool integrates Codon Usage analysis, providing real-time data on the translational impact of the modification. As shown in the output, the tool calculates the frequency difference between the original and modified codon (e.g., an AAG to AAA substitution resulting in a +1.4x usage increase), helping researchers avoid rare codons that could significantly decrease protein expression.

Finally, for direct one-step editing strategies, the “Generate Donor without PAM” module addresses the critical issue of Cas9 re-cleavage. The software identifies the PAM sequence associated with the input gRNA and proposes synonymous mutations to disrupt the recognition motif (Fig. 3G). This feature facilitates the design of “Hiding PAM” donors that prevent the nuclease from cutting the newly edited allele. Similar to the silent restriction site design, these suggestions are accompanied by a codon usage score, allowing the user to select the most translationally efficient variant that effectively masks the PAM, thereby maximizing the efficiency and stability of gene editing.

### Validating adaptive alleles using EasyCAPS

To demonstrate the practical utility of EasyCAPS, we applied the tool to assist the validation of quantitative trait nucleotides (QTNs) in the industrial-derived yeast strain PE-2_H4 [16]. Through our quantitative trait loci (QTL) mapping analysis of populations derived from crossing the PE-2_H4 strain with the S288C strain, we identified specific SNPs in the *HTA1*, *PHO84*, and *CAT5* genes associated with tolerance to lignocellulosic hydrolysate (LCH) [10]. We employed CRISPR-Cas9 allele swapping to transfer these favorable alleles into the laboratory strain, S288C. EasyCAPS was used to design strategies for “one-step” editing (Hiding PAM) and genotyping (Natural CAPS and dCAPS), streamlining the experimental validation (Fig. 4).

**Fig. 4.**
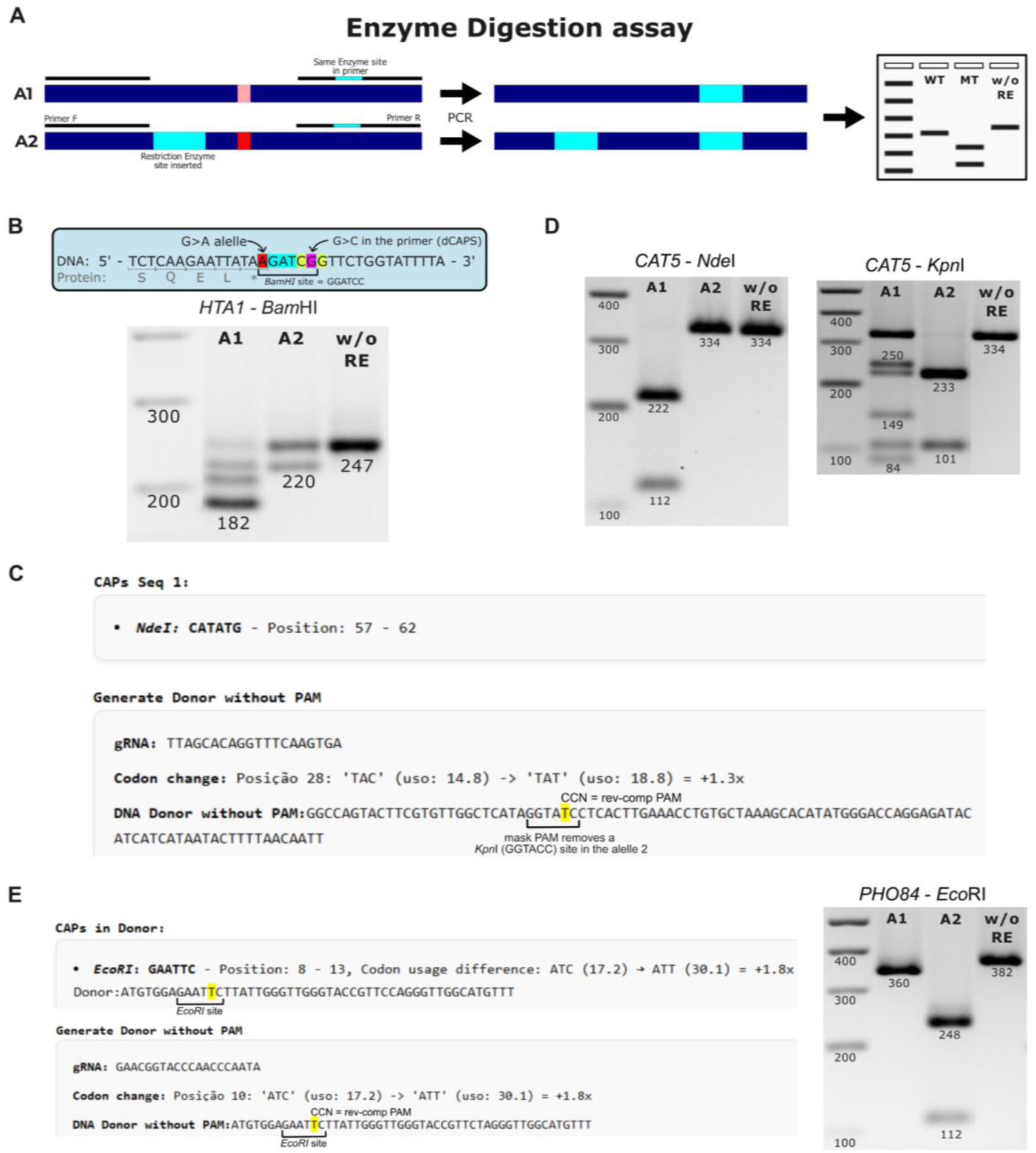
Experimental validation of CRISPR-Cas9-mediated allele swapping strategies facilitated by EasyCAPS. **(A)** Schematic of the enzyme digestion assay with an internal cleavage control. Primer R incorporates a restriction site (cyan box), ensuring a single cleavage in the allele 1 (A1) amplicon as a positive control. The edited allele (A2), containing an additional engineered site, undergoes double cleavage. **(B)** Manual design and restriction analysis for *HTA1*. A manually designed dCAPS primer introduces a mismatch to create an internal *Bam*HI site specifically with the wild-type allele, while the edited stop codon allele prevents this site formation. No PAM-masking mutation was included in this design. The digestion pattern of the 247 bp amplicon was partial; however, it still discriminates A1 (double cleavage; 180 bp major fragment) from A2 (single control cleavage; 220 bp). **(C)** EasyCAPS output for *CAT5*. The software identifies a natural *Nde*I site for genotyping and designs a PAM-masking mutation that removes a *Kpn*I site. **(D)** Dual restriction validation of *CAT5* editing. Left: *Nde*I cleaves A1 (222/112 bp) but leaves A2 intact (334 bp). Right: *Kpn*I cleaves A1 at two sites (149/101/84 bp), whereas A2 loses one site (233/101 bp). In the gel shown, the *Kpn*I digestion pattern in lane A1 was partial. **(E)** EasyCAPS donor design and validation for *PHO84*. Synonymous mutations were designed to generate a novel *Eco*RI site and disrupt the PAM. The adjacent gel confirms the edit via the internal control strategy (Panel A): *Eco*RI digestion of the 382 bp amplicon yields a single cleavage for A1 (360 bp) and a double cleavage for A2 (248/112 bp). **RS:** Restriction site; **RE:** Restriction enzyme; **w/o RE:** Undigested control.

To prevent false-negative results from incomplete digestion, we incorporated an internal cleavage control (Fig. 4A). One PCR primer includes the target restriction site, ensuring every amplicon undergoes at least one cleavage event. This confirms enzyme activity and differentiates alleles via single versus double cleavage patterns.

Initially, we targeted a stop-codon mutation in *HTA1* using a manual design prior to developing EasyCAPS. The strategy involved introducing a single point mutation at the stop codon to swap the alleles. To track this modification, a dCAPS primer was manually designed with a mismatch that, in combination with the wild-type sequence, artificially created a target *BamHI* restriction site. Notably, this manual design process focused strictly on genotyping and omitted a PAM-masking mutation, leaving the edited allele vulnerable to Cas9 re-cleavage. This omission exemplifies the complexities and pitfalls of manual experimental design, highlighting the specific need for the algorithms implemented in EasyCAPS. The assay successfully validated the allele swap: PCR amplification followed by *Bam*HI digestion discriminated the wild-type allele (A1), which formed the restriction site and underwent double cleavage (180 bp major fragment), from the edited allele (A2), where the stop-codon mutation prevented the *Bam*HI site formation, resulting in a single control cleavage (220 bp fragment) (Fig. 4B).

Subsequently, for the reconstruction of the T>C mutation in *PHO84*, we employed the full capabilities of EasyCAPS to facilitate a single-step editing strategy. The “Generate CAPS in Donor” module successfully designed a donor sequence containing a silent mutation that simultaneously removed the PAM site and introduced a novel *Eco*RI restriction site without significantly altering the amino acid sequence or codon usage (Fig. 4C). This automated design allows for the rapid screening of transformants. The *Eco*RI digestion of the PCR products yielded a differential pattern, producing fragments of 248 bp and 112 bp for the edited allele, distinct from the 360 bp wild-type fragment (Fig. 4D).

Finally, for the *CAT5* gene, we utilized the “Hiding PAM” and “Natural CAPS” modules to engineer a precise editing strategy. EasyCAPS identified a synonymous mutation in the PAM sequence with a favorable codon usage fold-change (+1.38x), ensuring efficient translation while preventing Cas9 re-cleavage (Fig. 4E). The successful integration of the donor was confirmed by dual restriction analysis predicted by the software: *Nde*I digestion tracked the targ*et all*ele (wild-type-specific cleavage, Fig. 4F - *Nde*I panel), while *Kpn*I digestion confirmed PAM modification (loss of a restriction site, Fig. 4F - *Kpn*I panel).

The successful installation of these alleles, validated using EasyCAPS-designed assays, enabled us to assess their biological impact through competition assays, revealing that the introduction of specific PE-2_H4 SNPs into the S288C background significantly improved fitness under LCH stress. Notably, the ease of tracking edited alleles with CAPS/dCAPS markers facilitated the rapid stacking of these adaptive SNPs with other beneficial alleles, resulting in cumulative increases in LCH tolerance in the engineered strains [10]. Together, these results illustrate how EasyCAPS streamlines molecular genetics workflows by bridging computational design and experimental validation in metabolic engineering and synthetic biology.

## Discussion

For decades, restriction enzyme-based genotyping has been used for SNP analysis, relying heavily on software for primer design. Pioneering tools such as dCAPS Finder 2.0 [3] and indCAPS [4] established the foundation for this technique; however, they were developed prior to the widespread availability of genomic data and were aimed mainly at plant molecular genetics applications. Users of these platforms often encounter significant technical limitations, such as strict limits on input sequence length, rigid, pre-defined lists of restriction enzymes, and outdated user interfaces that hamper high-throughput workflows. EasyCAPS directly addresses these bottlenecks by implementing a flexible architecture that accepts longer input sequences (up to 200 bp) and provides a dynamic and searchable enzyme library. This flexibility allows researchers to utilize enzymes currently available in their freezers, rather than ordering new reagents based on rigid software outputs. More importantly, EasyCAPS represents a methodological advancement by moving beyond the “genotyping-only” focus. While previous tools are limited to analyzing existing polymorphisms, EasyCAPS integrates dCAPS design with CRISPR-Cas9 engineering workflows. This capability allows users to simultaneously plan the introduction of a mutation (via donor design) and its subsequent validation (via CAPS/dCAPS), which is a very useful dual functionality.

Advancements in precision genome editing require tools that incorporate the biological context. In precision editing strategies, particularly those targeting essential genes or requiring “one-step” allelic replacement, preventing Cas9 re-cleavage is critical. Without automated guidance, manual design strategies might inadvertently mutate adjacent non-coding regulatory regions, such as 3’-UTRs, to abolish PAM sites, which is a risky approach that can unintentionally alter mRNA stability or gene expression [17]. Manually identifying the PAM sequence and designing a synonymous mutation that disrupts it without creating a new, unintended cut site or altering protein function is a multistep and error-prone process. EasyCAPS accelerates this phase by automating the “Hiding PAM” strategy. By rapidly suggesting valid synonymous mutations, this tool reduces the design time from hours to seconds. Furthermore, the integration of Codon Usage analysis distinguishes EasyCAPS from purely computational string-matching tools. Synonymous mutations are not biologically neutral; the introduction of rare codons can negatively impact mRNA stability and translation kinetics. By quantifying the fold-change in codon usage for every suggested mutation, EasyCAPS adds a layer of biological sophistication that safeguards protein expression levels, ensuring that the engineered strain remains physiologically relevant.

EasyCAPS integrates computational design and wet-lab validation. By streamlining complex tasks, such as the creation of silent restriction sites for donor tracking and the design of dCAPS primers for SNPs lacking natural sites, this tool simplifies the implementation of advanced genetic engineering. Its impact extends beyond the yeast models used in the present study. In plant breeding, where dCAPS markers are extensively used for marker-assisted selection (MAS) of agronomically important traits, EasyCAPS can facilitate the rapid development of markers for newly identified SNPs. In synthetic biology and metabolic engineering with microorganisms, this tool ensures the precise installation and traceability of pathway variants. Finally, in biomedical research, where CRISPR is used to generate cellular models of human genetic diseases, the ability to introduce specific patient alleles while efficiently masking PAM is highly advantageous. Ultimately, EasyCAPS empowers researchers to navigate the design-build-test cycle with greater speed, precision, and insight.

## Conclusion

In this study, we introduce EasyCAPS, a comprehensive web-based platform designed to streamline functional genomics workflows. By addressing the historical limitations of restriction-based genotyping software, specifically the constraints on sequence length and enzyme flexibility, EasyCAPS renews the utility of CAPS and dCAPS markers as cost-effective alternatives to sequencing for high-throughput screening.

Beyond genotyping, this tool fills a critical gap in the CRISPR-Cas9 editing landscape. The automated design of synonymous mutations for PAM masking (“Hiding PAM”) and donor engineering enables the execution of precise one-step allele-swapping strategies. This functionality effectively eliminates the risk of recurrent cleavage, a common bottleneck in homologous recombination-mediated editing, allowing efficient allelic conversion in a single transformation step. Furthermore, by integrating Codon Usage Bias analysis into the design process, EasyCAPS mitigates the biological risks associated with manual design, such as translational inefficiency or mRNA instability. Ultimately, EasyCAPS empowers researchers in diverse fields, from metabolic engineering to synthetic biology, to accelerate the design, build, and test cycle, making precise genome editing more accessible and robust.

## Declarations

### Consent for publication

Not applicable.

### Availability of data and materials

The EasyCAPS web application is freely accessible at https://easycaps-app.onrender.com/. The complete source code is open-source and deposited in the GitHub repository, available at https://github.com/olucasdebem/EasyCAPS_git.

### Competing interests

The authors declare no competing interests.

### Funding

This research was supported financially by the São Paulo Research Foundation (FAPESP) with grants 17/13972-1, 23/04162-7 to JG, and 17/24453-5, 22/14156-1 to APJ and 19/22263-0, 21/13906-4 and 23/06203-2 to LSB. A CNPq Universal project funding 404603/2023-8 was provided to JG.

## Acknowledgements

Not applicable.

## Notes

### Competing Interest Statement

The authors have declared no competing interest.

